# A simple demonstration of a privacy-preserving de-centralised genotype imputation workflow

**DOI:** 10.1101/2025.01.13.632689

**Authors:** Alban Letaillandier, David Picard-Druet, Thomas E. Ludwig, Gaëlle Marenne, Anthony F. Herzig

## Abstract

Recently, a number of studies have looked at the problem of privacy and data-sharing restrictions in the context of missing genotype imputation servers. This relates to the most typical imputation pipelines which involve a whole-genome sequenced haplotype reference panel being compared to genotyped study individuals (who have missing data to be imputed). Hence, involving two datasets from separate sources coming together in one informatic environment, where relatively complicated statistical models are applied; specifically, hidden Markov modelling. We give a short review of the current literature in this domain, observing three prevalent strategies: complicated data encryption, technical solutions to secure computation environments, and rearrangements of haplotype data in an effort to provide anonymisation. We embarked on a thought experiment to provide a potential fourth type of solution involving federating the different internal tasks within the statistical methods used for imputation. This idea is relevant considering there is currently motivation for federated analyses platforms in Europe for making combined inference across multiple genomic data resources. This allows for very simple manipulations to protect sensitive individual level data, which enable imputation algorithms to complete on simple plain-text files. We provide here an illustration of how such a federated imputation server could be put in place, along with associated code, including a simple implementation of the Li-Stephens haplotype mosaic model to achieve the imputation of missing genotypes. We name our general framework ANONYMP for anonymised imputation. A demonstration of the concept is given involving simulated data generated with msprime. We show that dividing different parts of the required calculations for statistical imputation between several sites is a valuable new avenue in the field of privacy-preserving imputation server development.

## Introduction

Despite the tumbling costs of whole-genome sequencing (WGS), the tried and tested strategy of array genotyping followed by haplotyping and missing genotype imputation still finds its place in modern genetic epidemiology study design [1,2]. WGS datasets come with multiple burdens: cost, data generation, storage, and computation capacity, and bioinformatic expertise. Imputation remains an attractive study design for many reasons. Firstly, the increasing number of human genomes that have been whole-genome sequenced has led to a greater number or increasingly large and diverse imputation panels being created [3–5]. There has also been an increase in the creation and availability of local reference panels [6–12]: panels of WGS datasets specific to different region of the global genetic landscape in human populations, often with a recruitment coming from a specific geographic region, cultural group but in most case from a specific nation. Secondly, there is an increasingly possibility to replace array-genotyping with low-coverage WGS [13–15]. This technique gives particular added benefits for imputation of individuals from parts of the ancestry spectrum for whom the choice of single-nucleotide polymorphisms (SNPs) for genotyping arrays is less pertinent for facilitating haplotype-matching. For the most part, genotyping arrays have been designed to capture population structure in European-ancestry populations [16,17].

An important consideration of imputation studies is that by nature it requires two datasets to work in harmony, which we will refer to as the reference panel and the target panel. The reference panel being the phased haplotypes from WGS for the reference individuals who are used for imputation and the target panel referring to the individuals with missing data, requiring imputation with array or low-coverage WGS data. In this work we only consider the scenario of array data for the target panel. In many circumstances, the two datasets are generated separately by different teams of researchers. Setting aside the multitude of potential difficulties in combing population genetic datasets without introducing any bias through batch effects, there is the added logistical challenge of finding an environment for the imputation to take place in a way such that the two owners of the two datasets remain within whatever regulatory and ethical constraints pertain to their data regarding the privacy of the participating individuals [18,19]. Typically, a haplotype reference panel represents a highly complex, versatile, and detailed resource; and hence a precious one. The combination of thousands of whole-genome sequencing samples (and hence containing many rare genetic variants), assigned to population labels and often with sample overlap between imputation panel and cohorts used in genome-wide association studies (GWAS) has unsurprisingly led to careful restrictions on the use of such panels. While certain panels are freely available, such as the combined call-set 1000G+HGDP [5] comprising the 1000 Genomes Project [20] individuals with those of the Human Genetic Diversity Panel [21,22], increasingly often they are only accessible via designated imputation servers, or require data-transfer agreements to be put in place specifically mentioning that such panels are only to be used for the purposes of imputation.

For many years, two possible imputation servers existed, Sanger [4] and Michigan [23]. It is at the Michigan server for example that the largest and generally regarded as most complete and highly performing imputation panel TOPMED [3] is available. Other imputation servers have recently gone online [24–29]. The idea of an imputation server is that it provides a safe environment for the reference panel to be housed and to avoid divulging the full dataset to each researcher wishing to perform imputation. Further imputation servers have gone online in other parts of the world, often to respond to the problem that researchers may have certain restrictions preventing them from sending their data to imputation servers. An example that the authors here are most familiar with being that GDPR (https://gdpr.eu/) prevents researcher in Europe sending out individual genetic data to the United States of America and hence the use of imputation servers outside of the EU is technically not permitted; and indeed the same issue would exist in the opposite direction [30] due to HIPAA guidelines in the United States of America.

This concern is not specific to the EU, there could be many circumstances where the two actors involved (the holder of the reference panel and the holder of the target panel) would have strict restrictions on the use and sharing of their datasets. This leads to a recent and niche research area of privacy-preserving methods for imputation servers, of which we give a short overview here. Our work serves as a commentary as well as putting forward potential additional methodological solutions to the logistical restriction of imputation servers.

A previous short review of privacy concerns and imputation was put forward by Sherman [31] which outlines the background of genotype imputation and imputation servers, current considerations regarding data-sharing that impact the potential usage of imputation servers and early work in the domain of privacy-preserving imputation. Three central themes appear in the current literature: 1) using complicated encryption methods to protect the data, 2) relying on a trusted secure third-party environment, and 3) manipulating reference panels to create ‘synthetic’ data such that imputation can still be achieved without directly using complete individual level haplotypes.

Kim et al. [32], Sakar et al. [33], and Gürsoy et al. [34] all approach the problem by proposing homomorphic encryption based calculations to achieve the imputation. Here, both reference and target data are safely encrypted and can be analysed together without de-encryption. The downside of this approach is an increase in computational burden coming from calculation on encrypted data and that neither method performs hidden Markov modelling of haplotype mosaics [35] for the imputation and hence have less precision than leading methods [23,36,37]. However, full implementations of HMM with homomorphic encryption are currently in development [38]. Dokmai et al. [39] demonstrate the loss of accuracy entailed by homomorphic encryption approaches and put forward an application of the minimac [23] software on Intel SGX’s Trusted Execution Environment framework; hence providing a solution relying on a highly trustworthy third party to perform the imputation. The same group have also recently extended their methods to provide a solution for haplotype phasing [40]. Such third-party server solutions have also been put in place for applications of genome-wide association studies (GWAS) [41–43]. Cavinato et al. [44] present a simple solution of creating new haplotype mosaics from a reference panel to render it anonymised and hence sharable. This avoids sharing complete individual level data though it would likely not prevent the re-identification of individuals that have contributed to the reference panel. The approach in question is elaborated on and developed further to also protect both the reference and target panel in Zhi et al. [45]. The key idea is that genotype imputation algorithms require haplotype data but only ‘locally’; to impute missing variants in a given genomic region, individual haplotypic data is required within and near that region but not across the whole genome. Hence, ‘mixing-up’ the haplotypes is not harmful to imputation. And even if haplotypes are broken and recomposed within the region in question, if this it done in a sensible way (with a realistic recombination model), the imputation algorithm can still deal well with this. As an aside, we also point the reader to Zhi et al. [45] for a thorough literature review of the current standpoints regarding privacy-preservation in the analysis of genomic data. Finally, Mosca and Cho [46] have recently evaluated the potential for the reconstruction of private genomes in the reference panel by carefully constructing synthetic target data; they also describe potential mitigation measures to avoid such attacks. A mention should also be made to Yelmen et al. [47] who use machine learning to create entirely synthetic imputation reference panels that are thus safe to share.

To summarize, the many recent papers on the subject show the interest of developing methodological solutions to allow for greater privacy preservation in the imputation-server framework. We observe the following: homomorphic encryption is appealing in nature but may be cumbersome to put in place in terms of additional calculation costs and potential loss of precision; going beyond two parties, so that the reference panel and target panel holders retain control of their sensitive data, and a trusted third party performs the imputation would seem essential; greater data protection can come from either breaking and reforming haplotypes [44], or simply by performing imputation in relatively small regions with different sample identifiers for reference and target haplotypes.

In this work, we present a combination of these different approaches, based on the following observations: first, a federated calculation environment would add data security by spreading across different nodes the different tasks involved in imputation algorithms, specifically in the hidden Markov models (HMMs); second, splitting the genome into many small regions would obscure the individual level data involved in both the reference and target panels, while losing little precision; and third, using simpler data encryption would facilitate calculations.

These approaches allow in a new potential anonymised imputation framework ANONYMP that we present here, to protect data using methods based on the following observations. The first key idea is that the Li-Stephens [35] HMM model used for imputation only needs the agreement status between reference panel and target panel alleles for all genetic positions present in the two datasets, as well as the distances between adjacent genetic positions. However, it is not essential to know where these positions are on the genome, nor what the alleles (nucleotides) are (they just need to be seen to be the same or not). Adding some noise in the genetic distance information is not significantly harmful to the HMM model accuracy and performance and helps hiding the genetic position. The second key idea is that the HMM runs only on the positions in common between the two panels, meaning that only these positions need to be shared with the external trusted parties for imputation. The third key idea is that while imputation works best with thousands of reference panel haplotypes available, typically far fewer are actually used when imputing a given target individual in a given region (see discussion of the K parameter later on in this work). The fourth key idea is that the output of the Li-Stephens HMM, posterior decoding via the forward-backward algorithm [48], do not need to be immediately used to calculate expected allele counts for missing genotypes in the target individuals, the output can be passed (with encryption) between actors.

The proposed framework remains largely a thought experiment: thus, we employ very naive ‘encryptions’ and a very simple application of the Li-Stephens model. Indeed, in this work we discuss encryptions in only a very loose sense. This work aims at assessing the feasibility of such a federated framework to be relevant for future cross-border projects involving federated calculation. This is an exploration of some new ideas into the emerging field of privacy-preserving imputation servers.

Methods

We present here a secure imputation framework involving five operators. The five include: the ‘1-User’, being the operator who holds the target-data to be imputed; the ‘2-Reference’, being the operator who holds the haplotype reference panel for imputation; and three external trusted third parties that we name ‘3-Compare’, ‘4-PPM’ (for posterior probability matrix), and ‘5-Product’. In practice, the last three external operators could potentially be combined in one centre with the possibility to have three different secure calculation nodes virtually isolated in hermetic informatic environments. The essential idea is that each operator only gets the data needed to realise a specific step of the imputation process. So, if the other security measures fail to prevent a single operator being compromised, the risk of leaked sensitive data is minimized. For this proof of concept, the ANONYMP implementation doesn’t focus on imputation results or performance. Furthermore, we don’t specify how operators communicate between each other, we just postulate that each operator can’t directly access files that others operators have not shared with them.

In Figure 1, we use the term ‘encryption’ in a loose sense, as we are simply performing some very basic modifications on the datasets. More strenuous encryptions could be made on the actual files and messages send between operators, but here we just focus on the simple encryptions or modifications to the files that render them non-informative in regards to individual level genotypes but nonetheless allow for HMM modelling.

**Figure 1.**
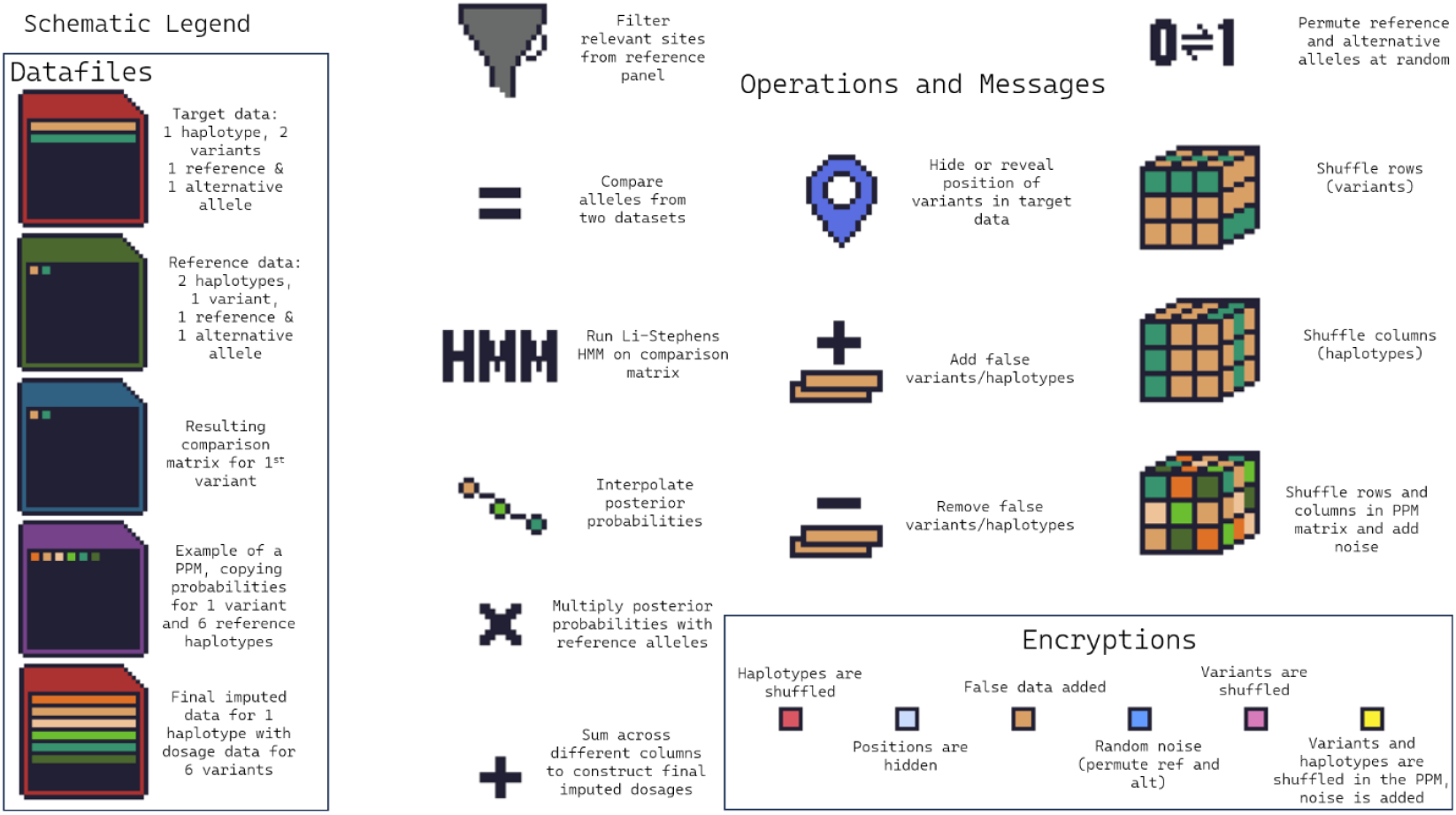
A detailed legend of the elements of the schematic in subsequent Figures (2 and 3) along with additional symbols for different types of possible simple random and reversible ‘encryptions’ including rearranging rows and columns, removing or hiding all chromosome and position information, permuting reference and alternative alleles in individual level data, adding spurious data, and adding random noise.

Figure 2 describes the role of each operator. 1-User provides the target genotype data to 3-Compare and get the final imputed genetic sequence from 5-Product at the end of the entire operation. 2-Reference provides subsets of the reference panel, firstly to 3-Compare and secondly to 5-Product. 3-Compare compares the target panel to the reference panel. It sends a comparison matrix to 4-PPM. 4-PPM computes the Posterior Probability Matrix (PPM, the output of the Li-Stephens HMM) from the comparison matrix and also perform linear interpolation of posterior probabilities. 5-Product computes the expected nucleotides for each SNP according to each haplotype. It does it by a simple product of the data from 2-Reference and the interpolated statistics provided by 4-PPM. This is then passed back to 1-User who sums this result over columns to form the final imputed dosages.

**Figure 2.**
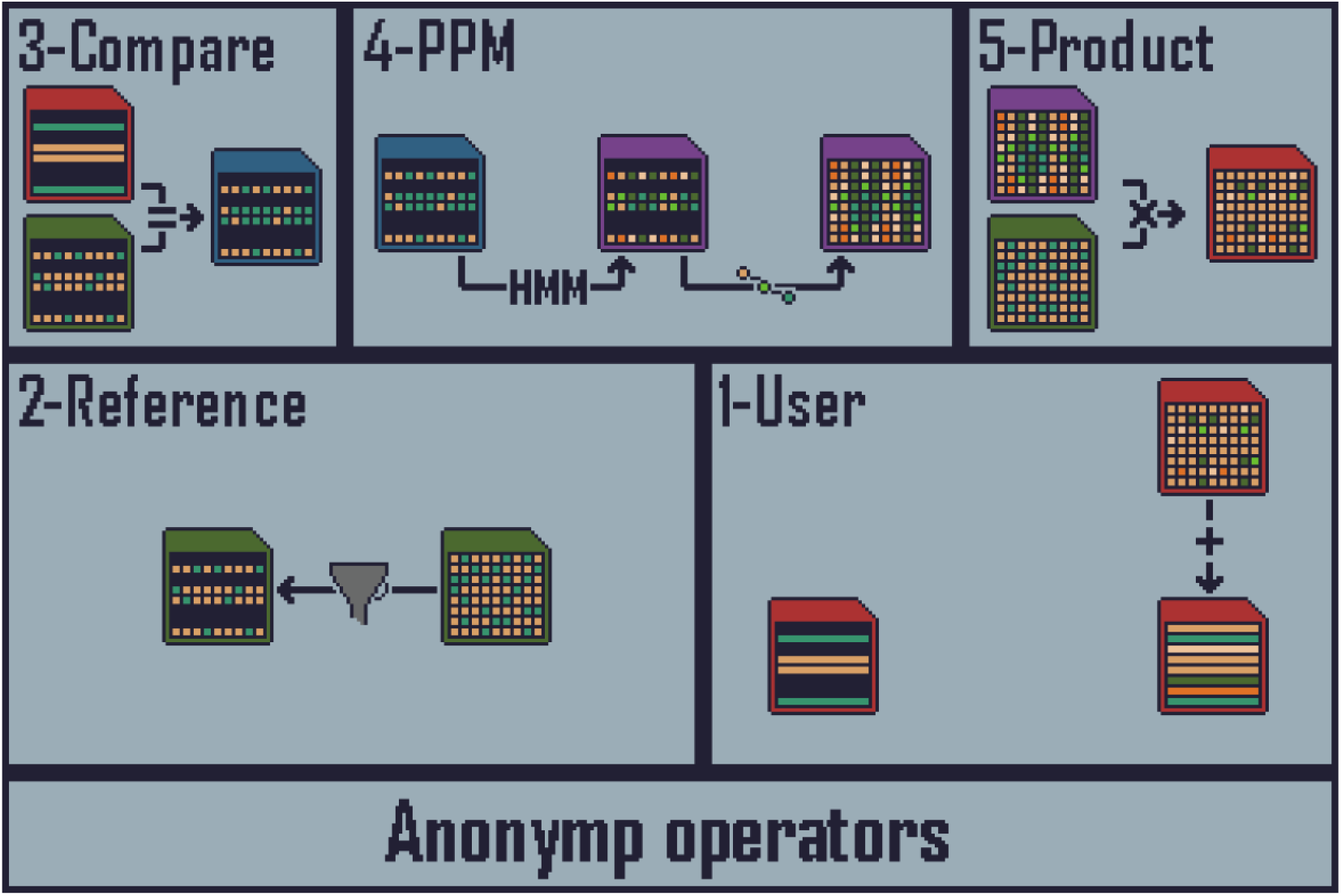
The different operators and operations are laid out. Datafiles are represented graphically with variants in rows and individuals in columns as described in (Figure 1. The schema starts in the bottom left corner of the 1-User box where we see a single target individual with genotyping array positions, target data is represented as red files, from there onwards the imputation protocol proceeds clockwise. The next step is to filter the same positions from the reference panel (green files in the 2-Reference box). Then both 1-User and 2-Reference sent their data to the 3-Compare server, where a comparison matrix (blue files) is generated which simply records the agreement or not of the two datasets. Agreement here refers to a logical test of whether two haplotypes carry the same allele at a given position. The comparison matrix is passed to the 4-PPM server where the HMM is performed to produce a purple posterior probability matrix, followed by linear interpolation to produce posterior probabilities at sequencing positions not present on the genotyping array. These posterior probabilities are then sent to 5-Product who also receives sequencing data of reference panel haplotypes from 2-Reference. 5-Product performs the required multiplications to calculate the component parts of the expected imputed minor allele counts for the target individuals. These components are then sent from 5-Product to 1-User, who combines them to for the final result: imputed dosages for previously missing genotypes.

In Figure 2, the operations that need to be performed are laid out and tasks are separated and allocated to the different operators. So far, no data security measures are added, for example 3-Compare has access to individual level data for both the target and reference panels. However, we could imagine that if the same encryption (especially something very simple) was added to both files, 3-Compare could still complete their tasks without accessing un-encrypted data. In fact, the comparison can be made with variants that are not in chromosomal order and could even involve completely spurious variants and haplotypes. The comparison also does not need any information about which chromosome or which variants are within the two files, just that they are the same set. The 4-PPM server requires this comparison matrix, that the variants are in order, that there are no spurious entries, and some (potentially rough) idea of genetic distances between adjacent variants in order to perform the HMM. The 5-Product operator needs to multiply linearly interpolated posterior copying probabilities with sequencing variants from the reference panel, again with no need for knowledge of chromosome position, and indeed random noise and spurious data points can be added to the calculation as long as they can be removed later.

To read through the summary given in Figure 3, begin at the bottom right corner of the 1-User panel. Then move along the bottom row of the 1-User panel, where first the information about chromosome and position are hidden, spurious data are added, all variants are coded as REF/ALT (denoting reference or alternative alleles) so no specific allele information is retained and the REF/ALT status of all alleles in the target haplotypes are permuted randomly, finally the order of the rows is randomised. Messages are exchanged with 2-Reference so that the exact same operations can be performed. Hence, when1-User and 2-Reference pass their modified files (denoted as file symbols with additional ‘encryptions’ represented by coloured squares on the top left corner of datafile symbols), they are encrypted in the same manner. 2-Reference makes an additional modification (bottom-right corner of the 2-Reference box); the order of the haplotypes in the reference panel are mixed up hence the 3-Compare server does not work with datafiles with the same column order for each iteration. The eagle-eyed reader may spot that this would all entail that the reference panel data is transferred a very large number of times to the 3-Compare server, as all these operations are specific to each target haplotype to be imputed. As this would be impractical, it may be more pragmatic if the 2-Reference operator were to send instructions to 3-Compare to re-shuffle and re-noise previously received data from previous imputation runs, in a manner to avoid ever ‘de-encrypting’ the reference data. This is entirely feasible if the encryptions proposed are ensured to be commutable operations and hence can be combined together to a single instruction involving changing 0s to 1s and vice-versa across the reference datasets (green files) held by 3-Compare. Hence, what is given in Figure 3 corresponds to the ‘first’ ever imputation run on the hypothetical multi-site platform. There are multiple ways to achieve the same goal, but the essential idea is that 3-Compare has two completely non-informative files but whose differences correspond to real differences between the alleles in the target and reference panel in a certain genomic region (note that 3-Compare does not need to know where this region is).

**Figure 3.**
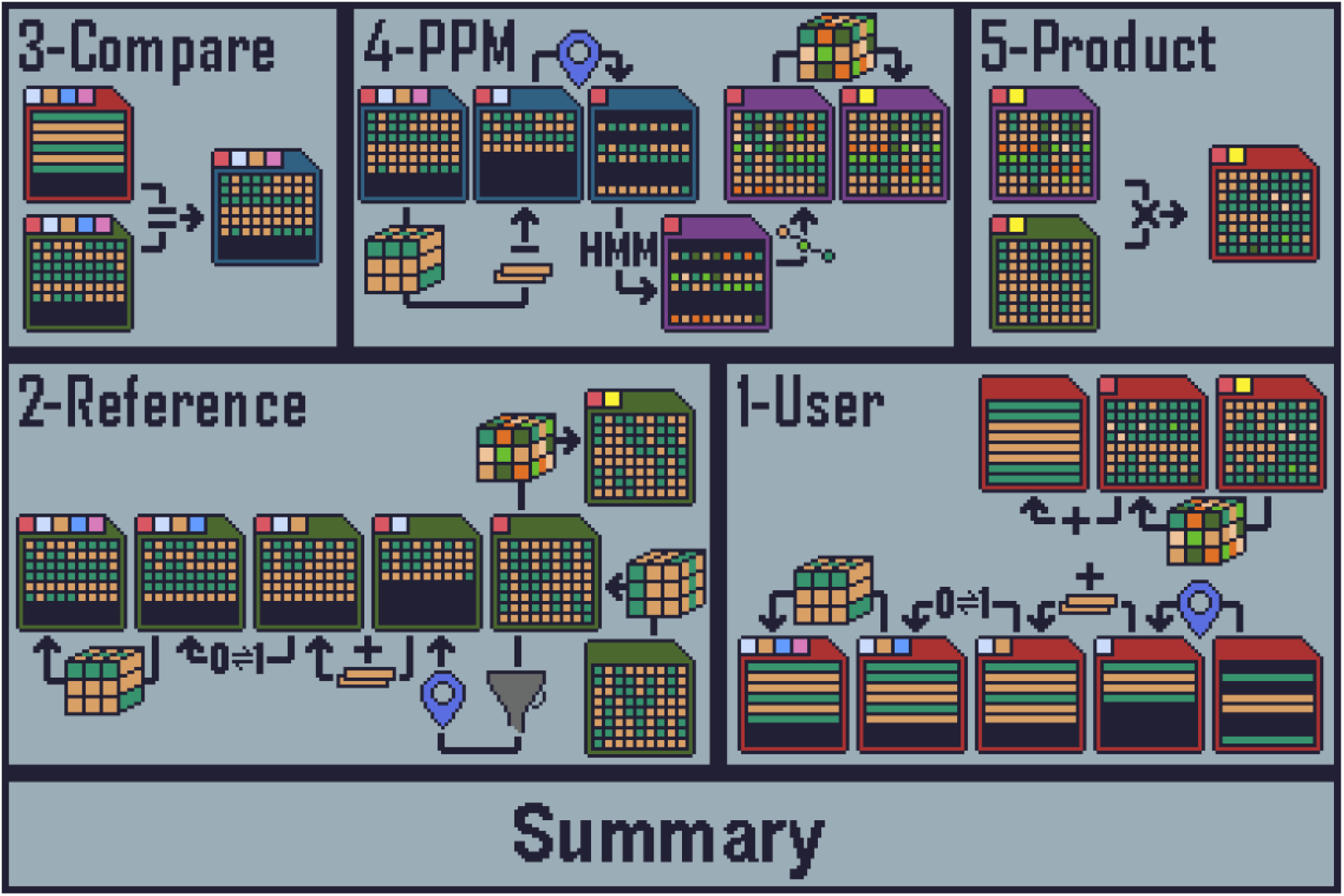
The full schematic of ANONYMP including the various data manipulations (detailed in Figure 1) to protect both reference and target data.

The comparison matrix (blue files) still potentially contains spurious entries, and the variants are not in genomic order, it hence not highly informative about the original data but some things can potentially be inferred. A row with mostly discordant data may well represent a variant which is likely rare and the target individual holds the alternative allele. This could begin to become dangerously informative once the variants (rows) are out back in order which is required to run the HMM. This is why we propose to separate this task and at this point pass to a 4^th^ operator: 4-PPM. Here, rows are put back in order, spurious entries are removed as 1-User has communicated with 4-PPM and revealed the required manipulations to do so. Even though there is no need to re-inject chromosome or position information, 4-PPM needs some estimation of recombination rates between adjacent sites, though this information can be very approximative without harming the imputation accuracy, which helps to avoid re-identification of the region based on comparison to a known genetic recombination map. The idea of adding noise to the recombination map is also discussed in Zhi et al. [45]. In our study, we either used the recombination maps provided by Bhérer at al. [49] or set a flat recombination rate between each pair of sites. At this point we would also recommend to anybody seeking to put in place an ANONYMP style system, to take advantage of the fact that most imputation software contain a ‘K’ parameter which details how many reference haplotypes are actually be considered for the forward-backward algorithm. This pre-selection of states can be performed by 3-Compare by simply inferring the hamming distance for each reference haplotype, simply by counting the number of discordant entries in the compare matrix in each column, and retaining only the K best scoring columns. This would essentially represent the pre-selection of reference haplotypes used by IMPUTE2 [50] and it is what we have put in place in the demonstrative code that we provide with this work. In this way, the 4-PPM server is only ever working with a very small subset of the reference panel, setting K=100 for example. Some precision in imputation could be lost at this point. Once 4-PPM has ran the HMM, the posterior probabilities need to be linearly interpolated to positions not present in the target data, but present in the reference panel data; requiring some communication with 2-Reference. Again, some noise could well be added regarding the exact positions of these reference panel only sites without any great harm to the imputation. At this point the final PPM matrix can then be completely reshuffled and noised by 4-PMM, and indeed need no longer be in matrix format, 2-Reference must do the same to their reference data (these steps are shown in the top-right corners of the boxes for 4-PPM and 2-Reference). At this point both 4-PPM and 2-Reference sent their shuffled and noised datafiles to 5-Product who simply multiplies the two together. 5-Product must be separate from 4-PPM as otherwise 4-PPM would be able to see the data of 2-Reference. The data must remain unordered and noised so that 5-Product does not see the final imputed data of the target haplotypes. Finally, 5-Product passes their output back to 1-User, who (after exchanging messages with 2-Reference) can finally re-order and denoise their data, and then sum across columns (this represents the weighted sums of different copying states from the HMM) to finally attain imputed genotype dosages. These final operations are shown in the top-right corner of the 1-User box.

In this way, only 1-User gets to see the final output of the imputation, and the reference and target data are always manipulated in encrypted formats. What we are describing here are very simple encryptions (rearranging data and adding noise) and hence all datafiles that need to manipulated remain in plain-text. However, more involved encryption such as discussed in the introduction could be used, we just aim here to put forward the possibility of avoiding such complexity.

## Results

To test ANONYMP, we took simulated data presented in [48] based on msprime [52] using the demographic scenario presented in [53]. Here we generated a reference panel of 45,000 individuals and a target dataset of 2000 individuals to be imputed. Code was developed to perform the ANONYMP imputation protocol and is made available with this work (see Data Availability Section). The five authors of the paper took on the roles of the five different operators (Figures 2 and 3) and were able to complete the imputation run; each working in a sperate working environment. The results were identical to an imputation run carried out directly; without the splitting of tasks and manipulations to data files.

Our simulated data only involved chromosome 15, which we split into 16 genomic regions or ‘chunks’ using scripts provided with IMPUTE5 [54]. In Figure 4 we compare the imputation accuracy of ANONYMP to IMPUTE5 and MINIMAC4 [23]. Our HMM likely performed a little worse than leading imputation software as our model was notably rather less elaborate in terms of selecting the K reference haplotypes to use for imputation or the estimation of model parameters for the HMM. The importance of the K parameter is apparent in Figure 4(a) and 4(d). However, it was not our goal to develop a highly efficient of high-performance imputation software but rather to provide a basic rendition of the Li-Stephens model to demonstrate the possibilities for federation and anonymisation during the imputation process. The extra task involved in manipulating the different datafiles add relatively little time compared to the actual HMM calculations required for the imputation as can be seen when running the example imputation runs as set out on the ANONYMP github. The HMM naturally takes the majority of the total time though it should be stressed that our implementation of the Li-Stephens was rather naïve, being just for the purposes of this thought experiment, and hence is orders of magnitude slower that the heavily optimised imputation software IMPUTE5 and MINIMAC4.

**Figure 4.**
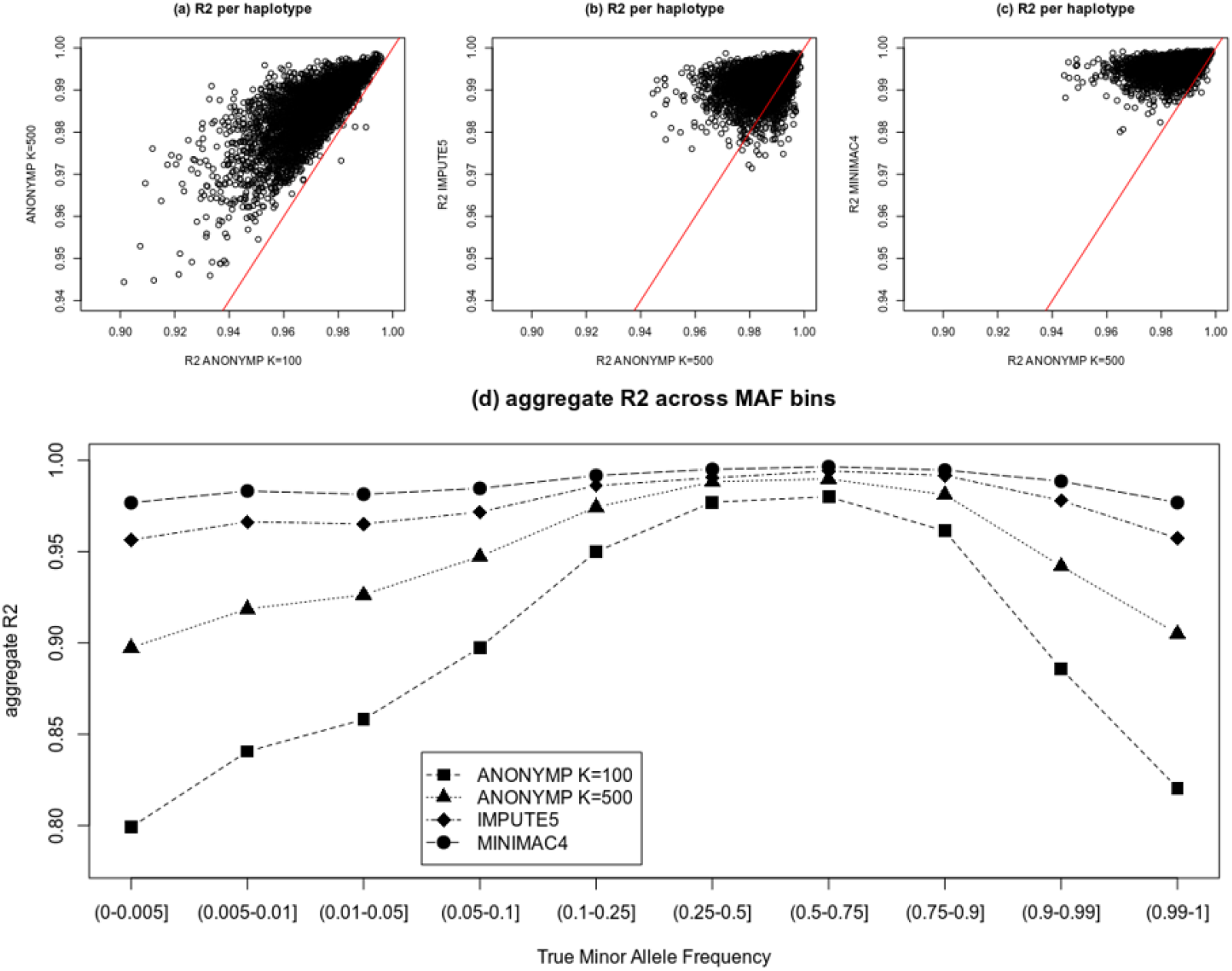
Comparison of imputation accuracy between ANONYMP, IMPUTE5 and MINIMAC4. In plots (a)-(c), squared correlations between imputed dosages and true allele are calculated across the 4000 haplotypes of the 2000 target individuals. We compare ANONYMP with two different settings: K=100 or K=500 where K describes the number of reference haplotypes used for the HMM. IMPUTE5 and MINIMAC were ran with default settings. In (d), aggregate correlations across all 2000 individuals were) calculated in different bins of minor allele frequency (MAF).

Finally, we assessed the ability of 4-PMM to attempt to guess the series of alleles for a given target haplotype by assigning a reference allele when the row in the comparison matrix contains a majority of agreements and an alternative allele otherwise. On average over the 4,000 haplotypes, and for K=100 and K=500, this simple method guesses only 63.3% and 68.4% of the alleles correctly, respectively. This was not much better than simply guessing all alleles as being reference alleles which gave an average of 62.8% correct allele guesses.

## Discussion

In this work, we have laid out some novel ideas for the development of privacy-preserving imputation servers that go beyond previously presented methods. The key innovation is to split or federate different elements of the imputation algorithm between three distinct calculation nodes (3-Compare, 4-PPM, and 5-Product). Our work offers some alternative strategies to previous studies, focusing less on complicated data encryption and secure environments, but on decentralising the analyses and using multiple computation nodes to split different internal tasks of the imputation models. The key contribution of this study is this innovation for improving strategies and workflows for privacy-preserving imputation servers. Our approach could easily be combined with other previously presented measures for protecting sensitive data during genotype imputation.

To explore the weaknesses of our strategy, we reasoned that the weakest link of ANONYMP would be at the 4-PPM server as here the variants within the genomic region to be imputed are put in order and genetic distances between adjacent cites may be given. 3-Compare and 5-Product have significantly less information. We showed that 4-PPM is unable to easily guess the target haplotype allele statuses (reference or alternative). But we should also consider the information that would be available for 4-PPM regarding the reference panel. In a given imputation run, the comparison matrix that has to be manipulated by 4-PPM resembles an MxK matrix of the K reference haplotypes selected by 3-Compare and the M variants present in both target and reference panels. The entries of the matrices are ‘true’ / ‘false’ values describing agreement between the target haplotype and each reference haplotype; information that might easily be stored as zeros and ones. This means that at this point 4-PPM essentially has a small portion of the haplotype reference panel; though without any point of reference regarding which rows or columns of the full haplotype reference panel it holds, or how the alleles are coded as here a zero is not necessarily aligned with a reference allele. Furthermore, this would allow 4-PPM to approximately calculate minorallele frequencies and linkage disequilibrium between sites. Combining this with genetic distances between adjacent sites (even if the distances have added noise) could conceivably begin to provide 4-PPM with enough tools to infer which part of the genome it is analysing; particularly if it can cross-reference with knowledge of the sets of sites that are present on commonly used genotyping arrays. We would argue that actually the majority of this information could also be gleamed during the calculation of forward and backward probabilities by a malicious actor. This would strengthen the argument for using a technique such a holomorphic encryption at the point of the 4-PPM calculation and also to ensure as much assurance regarding the trustworthiness of the environment where the imputation algorithm takes place; though in any case all efforts would be quickly undermined by collusion between actors who are not supposed to share information. We suggest that by splitting the calculation across different locations would therefore add extra protection.

At the sensitive 4-PPM stage, a few additional tricks could be added to aid in keeping this comparison matrix as uninformative as possible. By keeping K relatively low, leading to a small loss in accuracy, there would likely be a number of variants in full agreement across the K reference haplotypes with the target haplotype and these rows could be safely removed from the calculation by 3-Compare. There could also be variants in complete disagreement across the K reference haplotypes, which would correspond to rare alleles in the target haplotype. This eventuality is overall less likely but when occurring, again such rows could be removed (by 3-Compare) from the calculation with little impact. A more drastic approach would eventually be to drown out the possibility of 4-PPM making inference on the comparison matrices by simple asking it to perform more computation than necessary by mixing in additional synthetic imputation tasks and eventually even completely synthetic reference panels [47]. It would also probably be beneficial to split the chromosomes into sub-regions for imputation differently between different imputation runs; to avoid 4-PPM having the exact same dimensionality in the comparison matrices across all runs. As already discussed, at some point a more pragmatic approach would be to accept that at this point, if the HMM calculation is to be made correctly, it must involve either input and/or output with a certain similarity to the sensitive individual level data and that this cannot be avoided. In this case, 4-PPM would best be hosted on a very secure computational environment, a relevant example being the Helmholtz imputation server [27] or the environments presented in Dokmai et al. [39].

Two further final clear disadvantages of the ANONYMP framework are: firstly, the large number of messages and data-transfers that need to be made and secondly, that leading publicly available imputation software cannot be used as they are. The first consideration we predict could be overcome by changing slightly the schema proposed in Figure 3 so that for example instead of 2-Reference sending large files to 3-Compare, more simple instructions could be sent so that 3-Compare can re-arrange and re-noise previously received files from previous imputation jobs; similar ‘tricks’ could be easily applied in other moments in the schema to avoid repetitive large data transfers. Indeed, in the application of ANONYMP made available here, we specified that certain objects would always have a fixed size which both prevents certain operators guessing genomic locations and could facilitate such a recycling of previous messages of old imputation runs. In particular this would be relevant regarding the communication between 1-User and 2-Reference with 3-Compare, and the communications between 4-PPM and 5-Product and 2-Reference. For the second consideration, as we have shown, it is relatively simple to write a simple version of the Li-Stephens imputation algorithm but this was both slower and more inaccurate than current software which have undergone years of development and optimisation. Hence, if a federated imputation server with an ANONYMP-style solution would greatly benefit from collaboration with current imputation software developers in order to pick apart the different internal calculations and distribute them across different operators.

We conclude with a description of what was the essential motivation behind the thought experiment that led to developing ANONYMP; and why for the moment it remains largely a thought experiment. The 1+M Genome program was established by the European Commission in 2018 with an objective to align whole-genome sequencing efforts across the EU to provide inference from over 1 million high quality genomes for clinical and research purposes. In October 2024, a sub-project began named the Genome of Europe (https://b1mg-project.eu/1mg/genome-europe) with the objective of population-genetics analyses across 100,000 whole-genome sequences across 29 EU countries. One objective being to provide ancestry informed imputation haplotype reference panels with the caveat of each contributor to the project retaining their individual level WGS data within their own country. Analyses are to be federated between different calculation nodes as part of the Genomic Data Infrastructure (GDI) initiative. A plan is set out to collaborate with the recently opened Helmholtz imputation server where considerable work towards respecting GDPR considerations have been made. Given the multiplicity of potential calculation nodes via GDI and a secure third-party imputation environment in Munich could lead to a federated imputation approach, in the style of ANONYMP, being adopted. There would also be the important possibility to use such a set-up for more applications that imputation such as association testing [42] including association methods directly based on imputation algorithms [51]. But until a specific context (with its own specific constraints and objectives) where a privacy-preserving imputation server involving federated calculation is decided upon, we felt that it is not possible to specifically develop a new tool or federated imputation pipeline. Hence in this work we have simply laid out one potential type of solution, which either on its own or more likely in conjunction with other methodology for privacy-preserving imputation servers, could be a useful innovation in the field. We have made available all code developed during this project to facilitate future methodological developments and discourse on the subject of privacy-preserving imputation software.

## Data and software availability

All code and test datasets are available here: https://github.com/a-herzig/anonymp. Only simulated data were used in this study.

## Funding

This study was funded by INSERM under the Tremplin International First Step call (allocation number RSE24069NNA).

## Conflicts of Interest

The authors have nothing to declare

## Author contributions

The study was conceived by AL and AFH. Methodology was initially developed by AL and AFH and refined with discussion with all authors. Coding and implementation were performed by AL; analyses and simulation studies by AL and AFH. All authors contributed to testing, discussion and interpretation of results. The initial manuscript was written by AL and AFH, with all authors contributing to the final redaction. The study was supervised by AFH.

## Notes

### Competing Interest Statement

The authors have declared no competing interest.

